# Interaction of lecithin-cholesterol acyltransferase with lipid surfaces and apolipoprotein A-I derived peptides: implications for the cofactor mechanism of apolipoprotein A-I

**DOI:** 10.1101/225037

**Authors:** Marco G. Casteleijn, Petteri Parkkila, Tapani Viitala, Artturi Koivuniemi

## Abstract

Lecithin-cholesterol acyltransferase (LCAT) is an enzyme responsible for the formation of cholesteryl esters from cholesterol (CHOL) and phospholipid (PL) molecules in high-density lipoprotein (HDL) particles that play a crucial role in the reverse cholesterol transport and the development of coronary heart disease (CHD). However, it is poorly understood how LCAT interacts with lipoprotein surfaces and how apolipoprotein A-I (apoA-I) activates it. Thus, here we have studied the interactions between LCAT and lipids through extensive atomistic and coarse-grained molecular dynamics simulations to reveal mechanistic details behind the cholesterol esterification process catalyzed by LCAT. In addition, we studied the binding of LCAT to apoA-I derived peptides, and their effect on LCAT lipid association utilizing experimental surface sensitive biophysical methods. Our simulations show that LCAT anchors itself to lipoprotein surfaces by utilizing non-polar amino acids located in the membrane-binding domain and the active site tunnel opening. Meanwhile, the membrane anchoring hydrophobic amino acids attract cholesterol molecules next to them. The results also highlight the role of the lid-loop in the lipid binding and conformation of LCAT with respect to the lipid surface. The apoA-I derived peptides from the LCAT activating region bind to LCAT and promote its lipid surface interactions, although some of these peptides do not bind lipids individually. By means of free-energy calculations we provided a hypothetical explanation for this mechanism. We also found that the transfer free-energy of PL to the active site is consistent with the activation energy of LCAT. Furthermore, the entry of CHOL molecules into the active site becomes highly favorable by the acylation of SER181. The results provide substantial mechanistic insights concerning the activity of LCAT that may lead to the development of novel pharmacological agents preventing CHD in the future.

## INTRODUCTION

Coronary heart disease (CHD) and its comorbidities are global health threats showing an increased prevalence in industrial as well as in developing countries. Although many drugs for treating CHD exist, e.g. the cholesterol-lowering statins, a substantial residual vascular risk remains (Fruchart et al., 2014). According to the Residual Risk Reduction Initiative this represents a paramount public health challenge in the 21^st^ century (Fruchart et al., 2014). Thus, more effective medical strategies are needed to slow down and reverse this trend. It has been attributed that low high-density lipoprotein cholesterol (HDL-C) and elevated triglycerides levels are the two key contributors for the high residual CHD risk (Fruchart et al., 2008). Therefore, in the last decade, an increased focus has been on elevating the plasma concentration of HDL-C as a generally accepted intervention to prevent or reduce the development of CHD (Perk et al., 2012; Reiner, 2013). This opposed to the lowering of high low-density lipoprotein cholesterol (LDL-C), which in turn accelerates the fat accumulation into arterial walls. Traditionally, a high HDL-C concentration in blood has been regarded as a preventive measure reflecting the ability of HDL particles to transport cholesterol from the peripheral tissues back to the liver, including the mainly LDL-derived cholesterol from the arterial walls. This process is termed as reverse cholesterol transport (RCT) (Lewis and Rader, 2005; Tall, 1998). However, there is a growing body of literature showing that the elevation of the HDL-C by niacins or cholesterol ester transfer protein inhibitors does not or only moderately improve cardiovascular events (Group, 2017; Rader and Hovingh, 2014; Tall and Rader, 2017). In addition, recent genetic association studies have raised doubts towards the HDL-C hypothesis and causality of HDL-C in the development of CHD (Haase et al., 2012; Peloso et al., 2014; Rader and Hovingh, 2014). The failure of a large number of HDL-C raising therapies and the findings of genetic studies have led to a significant uncertainty concerning the benefit of raising HDL-C levels in the treatment of CHD. Still, several epidemiological studies support an inverse correlation between HDL-C and CHD (Després et al., 2000; Gordon et al., 1977; Rader and Hovingh, 2014). Hence, the in-depth understanding of the structure-function relationship of HDL particles and the enzymes processing them at the molecular level is all the more important for treating CHD.

The lecithin cholesterol acyl transferase (LCAT; E.C. 2.3.1.43) is one of the key enzymes driving the maturation of nascent HDL particles in serum (Jonas, 2000). Namely, LCAT is responsible for linking the acyl chain cleaved from a phospholipid to a cholesterol (CHOL) molecule resulting in a cholesterol ester (CE) molecule which in turn transforms discoidal HDL particles to spherical ones, which is a crucial step in RCT (Asztalos et al., 2007; Fielding and Fielding, 1995). Essentially, the esterification of CHOL molecules increases the cholesterol loading capacity of HDL particles enabling a more efficient RCT. In addition to its importance in RCT, mutations in the *LCAT* gene results in metabolic disorders such as familial LCAT deficiency and fish-eye disease in which the body's ability to metabolize cholesterol is severed, leading to corneal lipid deposition, haemolytic anaemia, accumulation of fatty material in arterial walls, and finally renal failure (Carlson and Philipson, 1979; Kuivenhoven et al., 1997, 1995).

The initial step in the reaction cycle of LCAT is its binding to a lipoprotein surface. Previous research studies have established that apolipoproteins, especially the principal LCAT activator apolipoprotein A-I (apoA-I), are playing a negligible role in the attachment rate of LCAT to lipoprotein surfaces, but decreases its detachment rate from surfaces (Adimoolam et al., 1998; Jin et al., 1999). This suggests that LCAT initially binds to the lipid moiety of lipoprotein particles which is then followed by interactions with apoA-I promoting stronger binding and activation. However, the specific amino acids responsible for anchoring LCAT initially to lipoprotein surfaces have not been revealed, although deletion mutant studies have shown that amino acids 53-71 and the disulphide bond formed between cysteines 50 and 74 of LCAT are crucial for the interfacial recognition of LCAT (Adimoolam and Jonas, 1997; Jin et al., 1999; Frank Peelman et al., 1999). Furthermore, it was shown by Murray et al. that when LCAT was bound to HDL or hydrophobic surfaces during sink immunoassays, the epitopes of the 121-136 region were not accessible for antibodies (Murray et al., 2001). Moreover, it is not known how deep the LCAT is buried into the lipid matrix, if a membrane binding region has specific interactions with different lipids that could facilitate its activity, and how LCAT is orientated with respect to lipoprotein surfaces when bound and not-bound to apoA-I. The next steps in the reaction cycle are the apoA-I driven activation of LCAT by an unknown mechanism and the diffusion of phospholipids (PLs) and cholesterol to the active site (Jonas, 2000; Sorci-Thomas et al., 2009). Consequently, the sn-2 chain of PLs is preferentially lipolyzed and the acyl-intermediate of LCAT is formed in which the acyl chain is covalently bound to SER181, which is a part of the ASP-HIS-SER catalytic triad found also in other lipases belonging to the α/β hydrolase family (Francone and Fielding, 1991; Jonas, 2000; F Peelman et al., 1999; Winkler et al., 199AD). Next, the acyl chain bound to SER181 is transferred to a free cholesterol. Finally, the newly synthesized CE molecule diffuses from the active site into the lipoprotein. While the LCAT activator region of apoA-I (central helixes 5, 6, and 7, or residues 121-142, 143-164. And 165-186, respectively) is roughly known based on the apoA-I LCAT deficiency mutations and vast amount of experimental data, the more specific LCAT interaction site and role of this in the different reaction steps remains unclear (Sorci-Thomas et al., 1993, 2009; Sorci-Thomas and Thomas, 2002). By revealing the mechanisms behind the different reaction steps of LCAT the way may be paved for inventing novel positive allosteric modulators of LCAT that aim to raise HDLC in a manner that would beneficial for the treatment of CHD.

For years, investigations concerning the LCAT lipid bilayer interaction and activation by apoA-I have been hindered by a lack of detailed atomistic structure of LCAT. Recently, however, X-ray structures of LCAT have become available enabling for example the computational studies of LCAT interacting with lipid surfaces and apoA-I to elucidate the mechanistic details concerning the cholesterol esterification process (Glukhova et al., 2015; Ruwanthi N Gunawardane et al., 2016; Manthei et al., 2017; Derek E. Piper et al., 2015). From the structures it is evident that the tertiary and secondary structure of LCAT is homological to lysosomal phospholipase A2 as pointed out by three similar folds: the cap-domain, the membrane-binding domain, and the α/β hydrolase domain (see Fig 1A) (Glukhova et al., 2015). Most importantly, structural details reveal that LCAT possesses a lid-loop that can move aside from the tunnel opening enabling the entry of lipids into the active site where the catalytic triad is located in, thus being consistent with other lipases (Ruwanthi N. Gunawardane et al., 2016; Derek E Piper et al., 2015).

**Figure 1.**
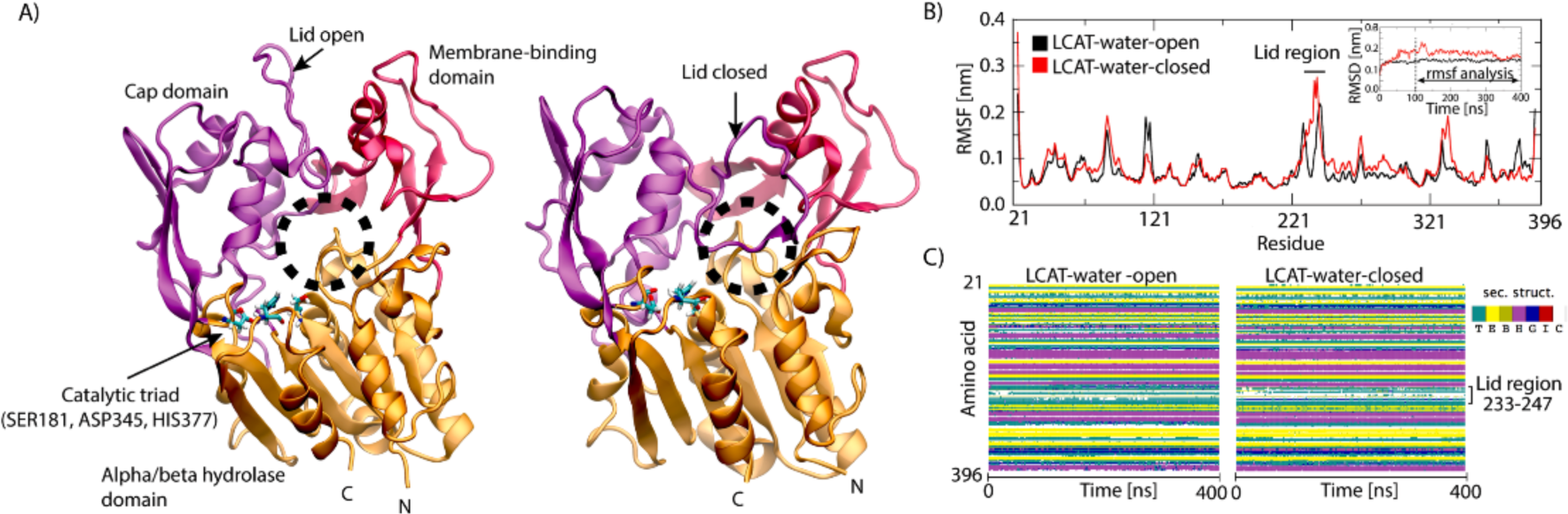
A) Structures of LCAT in the lid-open (left) and –closed (right) states rendered as cartoon models. The cap domain is coloured with purple, the α/β hydrolase domain with orange, and the membrane-binding domain with red. The tunnel opening of the active site is marked with a dashed black sphere. The catalytic triad residues: SER181, ASP345, and HIS377 are rendered with sticks and colored according to element types. Cyan atoms are carbon, blue nitrogen, white hydrogen, and red oxygen atoms. B) Root mean square fluctuation (RSMF) profiles for the lid-open and –closed states. Root mean square deviation (RMSD) results are shown in the inset. C) Secondary structure profiles for the lid-open and –closed states

In this study, we carried out several atomistic and coarse-grained molecular dynamics simulations to investigate the effect of different conformational states of LCAT to its interaction with a lipid bilayer comprised of dioleylphosphatidylcholines (DOPC) and CHOL molecules. To gain insights into apoA-I mediated LCAT activation, we utilized quartz-crystal-microbalance (QCM) and multi-parametric surface plasmon resonance (MP-SPR) experiments to investigate the effect of apoA-I derived peptides on the binding of LCAT to lipid bilayers, and the binding of LCAT to these peptides, respectively. These experiments were complemented with extensive free-energy simulations to reveal the role of the secondary structure of apoA-I derived peptides in lipid interactions. Finally, we studied the energetics of lipid ligand entry to the active site of LCAT.

Our simulations highlight the importance of specific non-polar amino acids in LCAT-lipid interactions. In addition, we show that the membrane anchoring non-polar amino acids attract CHOL molecules adjacent to them. The results also demonstrate that the lid-loop plays an important role in the conformation of LCAT with the respect of lipid surface. Furthermore, the experiments indicate that peptides derived from the LCAT activating region of apoA-I bind differently to LCAT and promote its lipid surface binding, although some of the peptides do not bind to lipids individually. We provide an explanation for this mechanism utilizing computational free-energy calculations. It was also found that the transfer free-energy of PL from the lipid bilayer to the active site is consistent with the activation energy of LCAT. Finally, our results indicate that the acyl-intermediate of LCAT highly facilitates the accessibility of CHOL molecules into the active site.

## RESULTS

### The flexible lid-loop covers non-polar amino acids located at the tunnel opening of LCAT from water in the closed state

To explore the structural and dynamical properties of the lid-loop region of LCAT, we carried out molecular dynamics simulations for LCAT in the lid open and closed states in water (Figure 1A). In this fashion, we strived to establish mechanistic insights to the conformational switching of the lid-loop that, presumably, is a prerequisite for the esterification reaction of LCAT to take place. Moreover, the overall stability of the LCAT structure was assessed against the X-ray structures to validate our computational models (Ruwanthi N. Gunawardane et al., 2016; Manthei et al., 2017; Derek E Piper et al., 2015). To construct applicable LCAT models for these purposes, the lid loop region (aa 233-247) was computationally grafted to the high resolution X-ray structures of LCAT representing either the open or closed state (Ruwanthi N. Gunawardane et al., 2016; Derek E. Piper et al., 2015). During this work, another high resolution X-ray structure of LCAT was published (PDB accession code: 5TXF) in the lid-closed conformation with only two amino acid residues missing from the lid-loop region (Manthei et al., 2017). This provided us an excellent means to validate our grafted LCAT model in the closed state. As shown in Figure S1 our model is in good agreement with the published LCAT structure in the closed state. The produced LCAT models were simulated up to 400 ns in water surroundings. These simulation systems were coined as LCAT-water-open-AA and LCAT-water-closed-AA systems (See more details in Material and Methods section).

First, the simulation trajectories were utilized to investigate the conformational stability, dynamics and secondary structure changes of LCAT. As it stands out from Figure 1A, the lid-loop exposes the active site for the entry of lipid ligands in its open state and shields the active site from water in the closed state. The root-mean-square deviation (RMSD) profiles in Fig 1B (inset) indicate that the backbone atoms of LCAT stabilized after 50 ns, and only small deviations from the initial structures occurred. This was reflected by the average RMSDs of 0.14-0.17 nm compared to the initial backbone atom scaffolds. By examining the mobility of LCAT as a function of residue number (RMSF profiles in Figure 1B), it was found that the lid-loop region (amino acids 233-247) shows the highest conformational fluctuations compared to the other structural parts of LCAT when neglecting the first few N- and C-terminal residues. This was an expected result. Concerning the secondary structure of LCAT (Figure 1C), the analysis showed a stable structure without significant changes during the simulations. A closer inspection revealed that the secondary structure of the lid-loop was a random-coil-rich in both conformational states, agreeing with the RMSF profiles that showed a high flexibility for this region.

Next, we analyzed the surface properties of LCAT and solvent accessible surface areas (SASAs) for the hydrophobic amino acids located at the active site tunnel opening (Figure 2A) since we hypothesized that the lid-loop shields these amino acids before LCAT detaches from the lipoprotein surface to the water phase. As shown in Figure 2B, the surface representations of LCAT as a function of residue types (non-polar, polar, and charged) reveal that hydrophobic amino acids located at the tunnel opening of the active site are exposed to the water phase in the open state and shielded from the water in the closed state. This result was supported by SASA calculations (averages from the last 300 ns) indicating that four of five hydrophobic residues located at the active site tunnel opening became shielded from the water in the closed state (Figure 2C). More specifically, these amino acids were LEU62, PHE67, LEU117, and ALA118.

**Figure 2.**
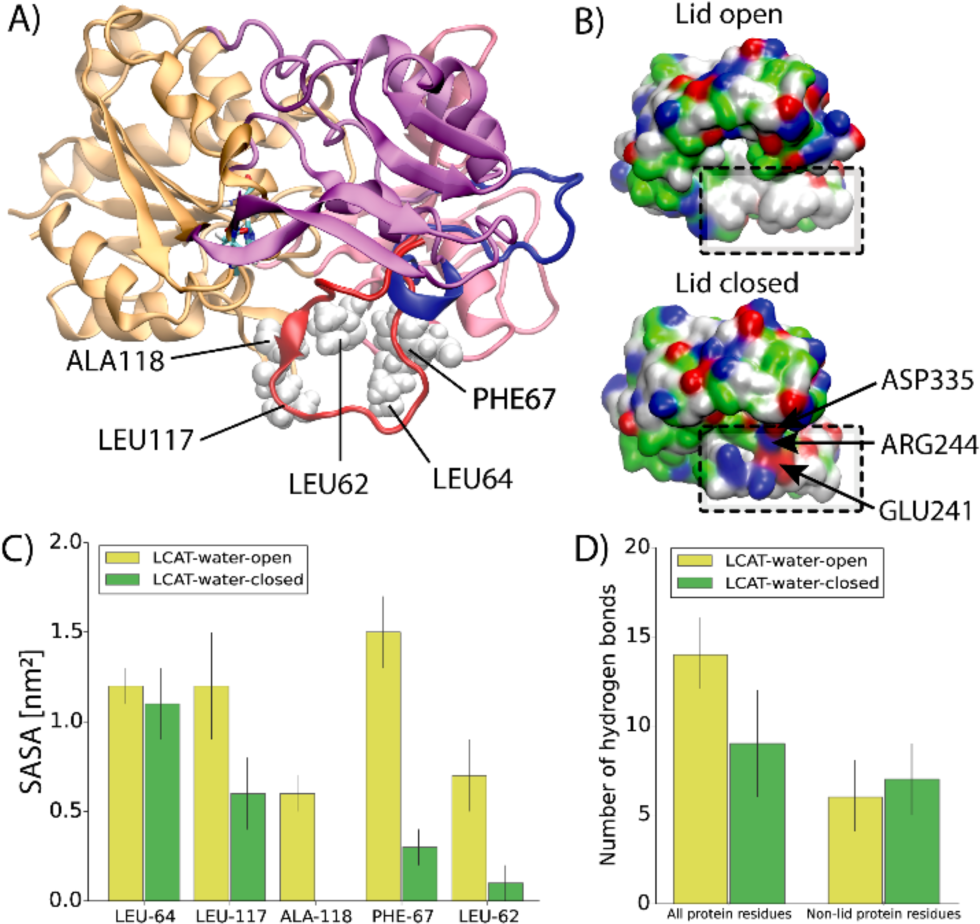
A) Structure of LCAT showing the hydrophobic amino acids located at the tunnel opening of the active site. LCAT is rendered with cartoon representation, and the domains are colored as in Fig 1. Additionally, the lid region of LCAT is colored with blue (the open state) or red (the closed state). Non-polar amino acids interacting with the lid loop in the closed state are labelled with the corresponding residue names and rendered with white Van der Waals spheres. B) The surface presentations for LCAT in the lid-open and lid-closed conformations. The location of the hydrophobic region is marked with black and dashed squares. Red indicates negatively charged, blue positively charged, green polar, and white hydrophobic amino acid residues. The salt-bridges formed by the lid-loop are marked with arrows. C) The average solvent accessible surface areas (SASAs) for the selected non-polar amino acids in the lid-open and -closed states. D) The average number of hydrogen bonds between the lid loop and all protein residues or non-lid protein residues in the lid-open and –closed states. Errors bars are standard deviations

Considering the previous finding that hydrogen bonds and salt-bridges play a role in the lidloop conformational switching of pancreatic lipase (van Tilbeurgh et al., 1992), we also evaluated the average number of hydrogen bonds and salt bridges formed in the lid-closed and -open states. Firstly, we calculated the average number of hydrogen bonds formed between the lid-loop and all LCAT amino acids. Secondly, we analyzed the average number of hydrogen bonds formed between the lid-loop and the non-lid-loop amino acids of LCAT. The results revealed that the lid-loop forms five internal hydrogen bonds when the conformational change from the closed to the open state takes place (Figure 2D). However, the salt-bridge analysis indicated the breakage of two salt-bridges formed by ARG244-ASP335 and ARG244-GLU241 at the same time (Figure 2B). Our analysis did not register stable salt-bridges between the lid-loop and the non-lid-loop residues in the lid-open state. In addition, no internally formed lid-loop salt-bridges were found in the open state.

### The lid-open conformation enables a deeper burial of the tunnel opening non-polar amino acids of LCAT at lipid surfaces

In the previous section, we established that LCAT exposes the non-polar amino acids located at the tunnel opening to water while the lid is open. Next, we asked if these non-polar amino acids could also interact with lipid bilayers resembling the lipid moiety of discoidal HDL particles. Additionally, we hypothesized that the lid-closed conformation also prevents the burial of the tunnel opening non-polar amino acids to a lipid matrix, which might be a prerequisite for the entry of lipid ligands in addition to the lid-open state. Therefore, we carried out atomistic and coarse-grained molecular dynamics simulations of LCAT to investigate how these non-polar amino acids interact with a bilayer composed of dioleoylphosphatidylcholines (DOPC) and cholesterol (CHOL). LCAT was initially placed on the surface of an equilibrated bilayer in a way which enabled the interaction of the tunnel opening non-polar residues with the lipid matrix. Both the lid-closed and -open states of LCAT were modelled. The atomistic simulations (termed as LCAT-mem-open-AA and LCAT-mem-closed-AA) enabled us to study the atom-scale interactions between LCAT and individual lipids in a microsecond time window. On the other hand, the coarse-grained representations (LCAT-mem-open-CG and LCAT-mem-closed-CG) extended the time window up to several microseconds rendering it possible to estimate e.g. the free-energy of binding of LCAT to a lipid bilayer in lid-open and –closed states utilizing umbrella sampling simulations.

Simulation trajectories revealed that LCAT stays at the surface of lipid bilayer up to 1 μs or 20 μs in atomistic and coarse-grained simulations, respectively (see Movies S1-S4). What stands out from the trajectories is that the non-polar amino acids of the tunnel opening and the membrane-binding region stay buried in the lipid matrix in LCAT-mem-open-AA and LCAT-mem-open-CG simulations (Figure 3A and Movies S1-S2). However, the lid-closed conformation prevents the deeper burial of tunnel opening non-polar amino acids to the lipid matrix (Movies S3-S4). Closer inspection by utilizing distance calculations with respect to the phosphorous atoms of DOPCs, revealed that the hydrophobic amino acids: L117, L64, F67, W48, L68, and L70 were clearly located in the acyl chain region of phospholipids in LCAT-mem-open-AA simulation (see the left panel in Fig 3B). From the distance analysis, we can see that these amino acids became less buried in the lipids in the lid-closed conformation, especially amino acids A118, L117, L64, F67, W48, L68, and L70 (see also Movie S3). As shown in Figure 3B, this trend was also seen in CG-simulations further verifying our initial hypothesis that the lid configuration plays an important role in regulating the interaction mode of LCAT with lipids. Further analysis revealed that the lid-closed state also changes the orientation of LCAT with respect to the lipid bilayer surface that explains the decreased burial of A118, L117, and L64 located at the hydrophobic tunnel opening. This is shown by a higher average tilt angle of LCAT calculated between the vector, formed by the C_α_-atoms of MET49 and ASN131, and the lipid bilayer normal (Figure 3B and Movies S2-S4).

**Figure 3.**
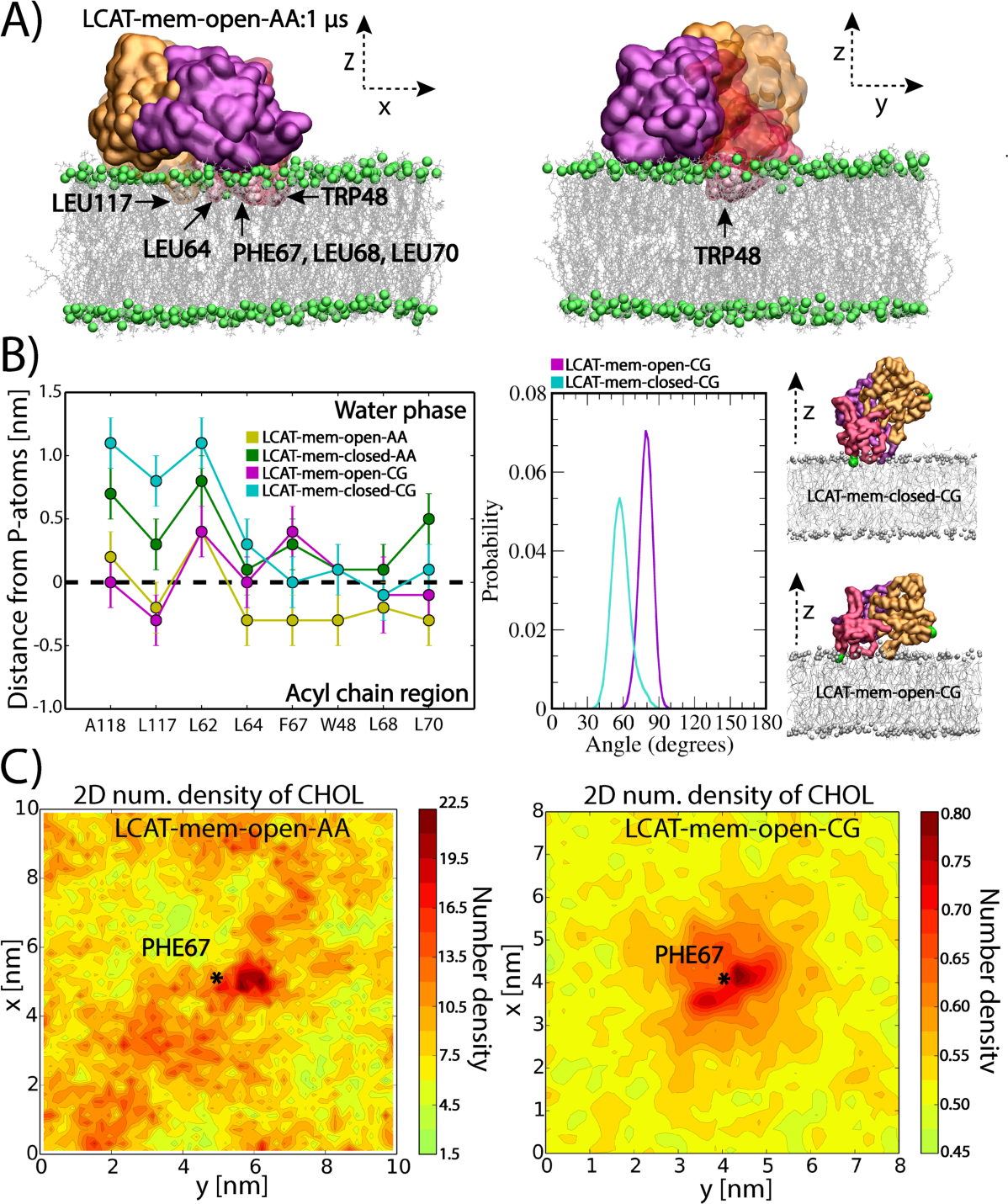
A) Snapshots from the end of LCAT-mem-open-AA simulation (1μs). The coloring of LCAT domains is following: Orange is the alpha/beta-hydrolase, purple the cap, and red the membrane-binding domain. Grey sticks represent DOPC molecules and green spheres phosphorous atoms of DOPC. Water molecules have been removed from the snapshots for clarity. The membrane penetrating hydrophobic residues of LCAT are marked and labelled to the snapshots. B) The average center of mass distances from the phosphorous atoms of DOPCs for the lipid-buried non-polar amino acids in the lid-open and lid-closed membrane simulations (left). The average tilt angle of LCAT with respect to the normal of lipid membrane in LCAT-mem-open-CG and LCAT-mem-closed-CG simulations (right). In addition, snapshots approximating the average tilt angle in each case are shown. Coloring is the same as in panel A, but the tilt vector forming amino acids (ASN131 and MET49) have been marked with green spheres. C) The 2D-number density maps for LCAT-mem-open-AA and LCAT-mem-open-CG simulations. The center of mass of PHE67 is marked by a star showing the location of the membrane penetrating region of LCAT

Next, we asked if the tunnel opening and membrane anchoring regions can interact with DOPC or CHOL molecules in a more specific way. To characterize this, we analyzed the 2D-number densities by first dividing the XY-plane into 100x100 or 36x36 squares depending on the system studied, atomistic or coarse-grained representation, respectively. This was followed by the calculation of the number densities of DOPC and CHOL atoms in each square (See more details in Materials and Methods section). We found no specific binding in the case of DOPC molecules, but CHOL molecules preferred to accumulate next to the membrane penetrating region of LCAT in both atomistic and coarse-grained simulations (Figure 3C and Figure S2).

### LCAT-interaction region derived peptides of ApoA-I facilitate the binding of LCAT to a lipid surface

To gain additional mechanistic insights regarding the interactions of LCAT with lipids and the possible role of apoA-I in this, we studied the binding of LCAT to a lipid bilayer with and without apoA-I derived peptides by employing the quartz crystal microbalance (QCM) technique. The apoA-I derived peptides were selected from the previously proposed LCAT-activation region (Sorci-Thomas et al., 2009) and were based on the apoA-I amino acids of 122-142, 135-155, and 150-170 (Figure 4A). In addition, a control peptide was chosen from the region of apoA-I comprising amino acids 185-205 which is not involved in the activation of LCAT. The effect of mutation Y166F on the properties of peptide 150-170 was also studied since it has been demonstrated that this mutation hampers the activity of LCAT (Wu et al., 2007).

**Figure 4.**
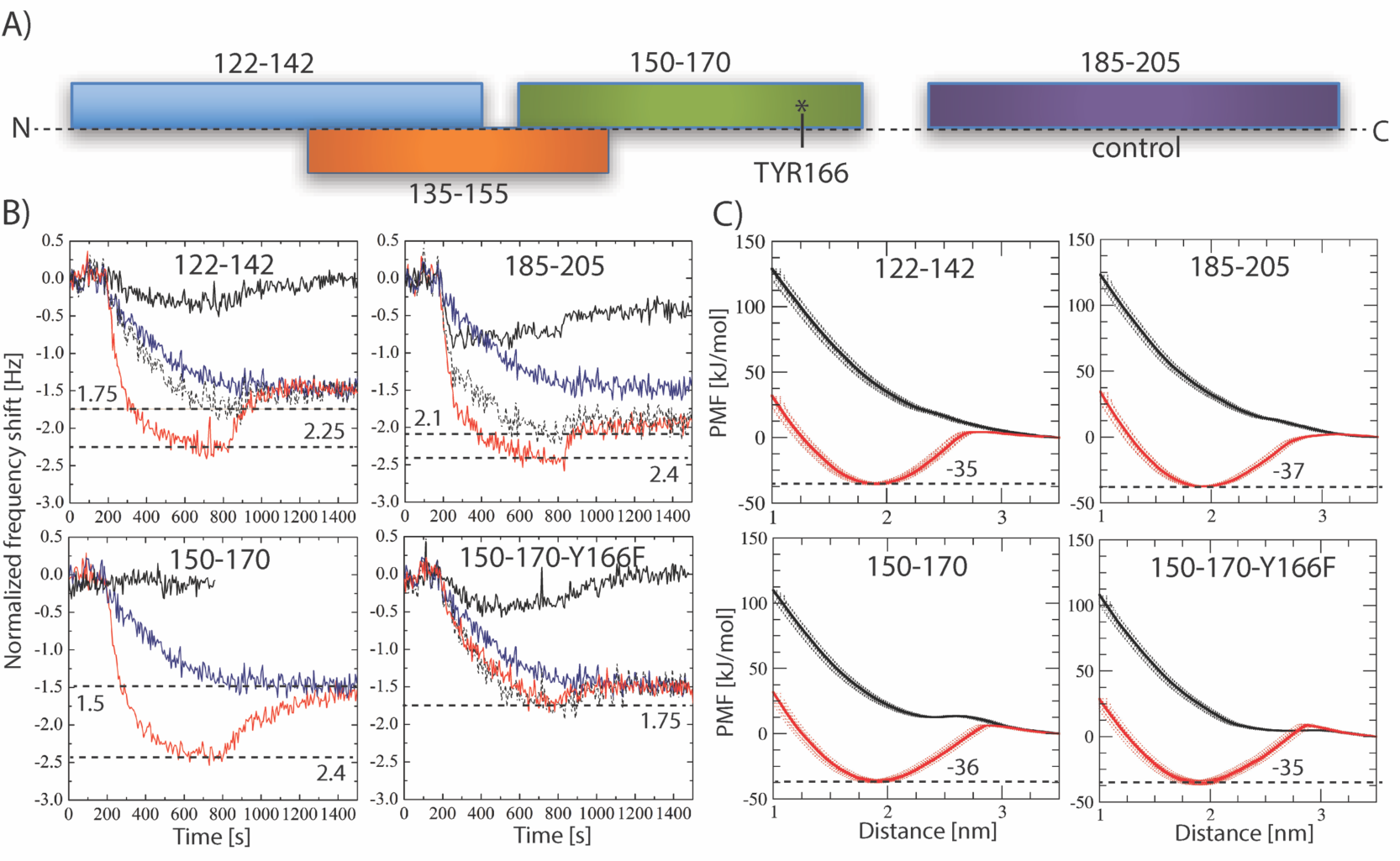
A) Schematic illustration showing the peptides studied along the sequence of apolipoprotein A-I B) Quartz crystal microbalance (QCM) responses for the pure LCAT (blue curve) and peptides (black curves) on the surface of the POPC bilayer. Surface responses of LCAT in conjunction with peptides are shown in red. The sum curve of pure LCAT and peptide measurements is marked as a dashed black curve. The horizontal dashed lines mark the minimum responses for different measurements, including their value. The result for peptide 135-155 is not shown, but the profile is similar as in the case of peptide 150-170 (See SI). C) Potential mean force (PMF) calculations for different peptides in coil (black) or helical (red) secondary structure as a function of distance from the center of a lipid bilayer. The binding free-energies for the peptides (kJ/mol) are marked next to the dashed black lines

The QCM results in Fig 4B indicate that LCAT binds to the POPC membrane without apoA-I agreeing with previous experimental studies and our simulation results showing that LCAT stays at the lipid membrane utilizing the tunnel opening non-polar amino acids or membrane-binding region in the attachment. The results also pointed out that some of the apoA-I derived peptides did not interact with lipids, namely only peptides 122-142, 185-205, and 150-170-Y166F were found to bind to the lipid bilayer (See Fig4B, solid black curve). The combined interaction studies of LCAT and peptides revealed that peptide regions 122-142, 135-155, and 150-170 clearly increased the affinity of LCAT to the lipid bilayer (Figure 4B). Results also showed that peptides 185-205 and 150-170-Y166F did not increase the lipid bilayer affinity of LCAT.

As peptides 150-170 and 135-155 did not interact with the lipid surface individually, but still increased the overall binding of LCAT, we hypothesized that the putative apoA-I interaction site of LCAT could induce amphipathic α-helical structures for the peptides driving stronger lipid interaction due to increased lipophilicity of the LCAT-peptide complexes. Another hypothesis considered the possible role of the peptides in changing the conformation of the lid from the closed to the open state enabling a stronger interaction of LCAT with lipids.

To test the first hypothesis, we calculated the potential mean force profiles (PMFs) for each peptide when they are transferred from the water phase to the lipid-water interface utilizing CG-simulations. We carried out calculations with both coil and α-helix secondary structures to investigate the role of the secondary structure of apoA-I derived peptides regarding the membrane interactions. The results depicted in Figure 4C indicate that all peptides can bind to the surface of the lipid bilayer if they adopt α-helix conformation. However, none of the peptides showed a free-energy minimum within the membrane region in the coil form. Therefore, for a peptide region to induce stronger binding of LCAT to the lipid membrane surface they must fully or partly adopt an amphipathic α-helical secondary structure to interact with the unknown apoA-I interface of LCAT.

To assess the possibility of the second hypothesis, we calculated the binding free-energies for LCAT in open and closed states. As shown by the PMF profiles in Figure 5A, the lid-open state (ΔG_water->lipid_ = −28±1 kJ/mol) is not able to bind so strongly to lipids when compared to the lid-closed state (ΔG_water->lipid_ = −34±1 kJ/mol), although the previous findings indicated a deeper lipid interaction for the open-state (Figure 3B). The value for the closed state is well in agreement with existing experimentally determined dissociation constants giving binding a free-energy value of −36 kJ/mol for LCAT when interacting with small unilamellar vesicles in the absence of apolipoproteins (Jin et al., 1999).

**Figure 5.**
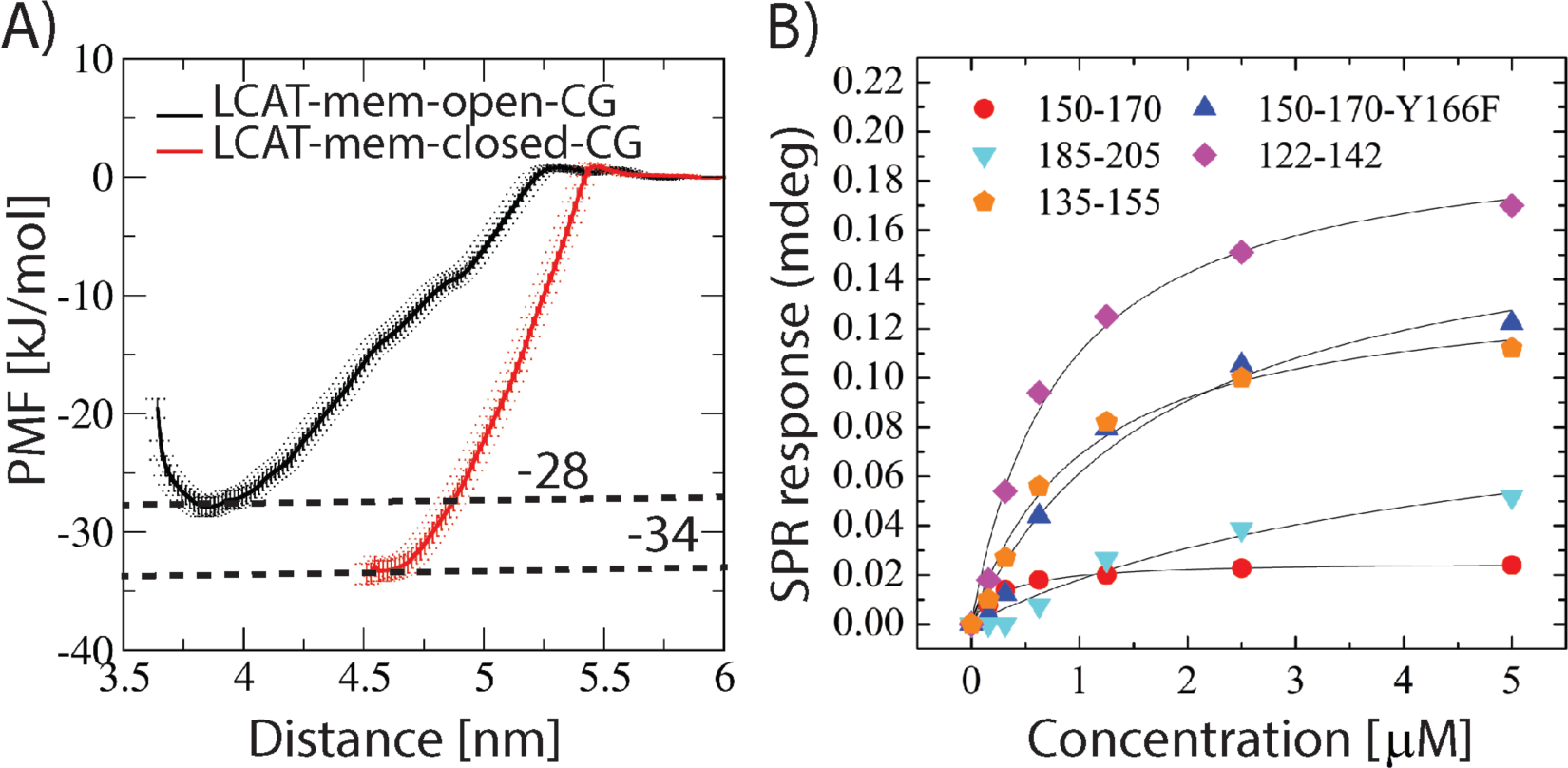
A) Potential mean force profiles for LCAT as a function of distance from the center of lipid bilayer. The transfer free-energies of LCAT from water to the lipid bilayer surface are marked next to the dashed lines (kJ/mol). B) Surface plasmon resonance peak responses for different apolipoprotein A-I derived peptides as a function of LCAT concentration

To also elucidate the specificity of different peptide-regions against LCAT, we carried out multi-parametric surface plasmon resonance (MP-SPR) experiments to determine dissociation constants (K_D_) for the LCAT-peptide complexes. The peptides were attached to streptavidin-coated gold-sensors via a biotin anchor and LCAT was flowed over the anchored peptides to measure the K_D_:s using a Langmuir model (See more details in Material and Methods). From the MP-SPR results (Figure 5B and Table 1) it can be deduced that the three strongest affinities of LCAT were with peptide-regions 150-170, 122-142, and 135-155, while the lowest affinities were against peptides 185-205 (control) and 150-170-Y166F. These results suggest that LCAT possesses a more specific binding site for peptides 150-170, 122-142, and 135-155. This is also corroborated by the QCM results which correlate well with the affinity values obtained from SPR measurements (Table 1).

**Table 1.** MP-SPR derived dissociation constants (K_d_) and R_max_ values for different LCAT-peptide complexes. In addition, the QCM response differences between the combined LCAT+peptide and the sum of separate (LCAT and peptide alone) measurements are given (QCM-diff.).

| Peptide | $K_d$ (μM) | $R_{max}$ | QCM-diff. |
| --- | --- | --- | --- |
| 122-142 | 1.0258 | 139.1 | -0.5 |
| 135-155 | 0.8193 | 301.0 | -1.0 |
| 150-170 | 0.2903 | 25.4 | -0.9 |
| 150-170-Y177F | 1.8442 | 174.4 | 0 |
| 185-205 | 4.6901 | 103.4 | -0.3 |

### The acyl-intermediate of LCAT lures cholesterol molecules to the active site

The entry of lipids to the active site of LCAT is still mechanistically an unknown process (Jonas, 2000; Sorci-Thomas et al., 2009). Especially, the possible direct role of apolipoproteins is unknown. To gain additional insight into this matter, we calculated potential mean force (PMF) profiles for DOPC and CHOL molecules when they were pulled from the lipid matrix to the active site of LCAT. PMF calculations were carried out utilizing the equilibrated conformation of LCAT in the LCAT-mem-open-AA simulation. We assumed that the lid should be in the open state before entry of ligands.

The results in Figure 6 indicate that the free-energy costs of pulling DOPC or CHOL molecule from the lipid matrix to the active site are approximately 65±2kJ/mol and 35±4 kJ/mol, respectively. The transfer free-energy cost of DOPC matched well with the experimental LCAT activation energies that range from 53 to 76 kJ/mol measured for HDL-LCAT complexes comprised of POPC or DOPC lipids (Jonas, 1986; Parks and Gebre, 1997). Next, we asked if the acylation of SER181 decreases the free-energy cost needed for CHOL to enter the active site. Thus, we constructed an atomistic simulation system where SER181 was acylated and the lid was in the open state (LCAT-mem-acyl-AA). Surprisingly, we found that the acyl-intermediate of LCAT did not only decrease the free-energy cost but rendered the entry of CHOL to the active site almost favorable reflected by the transfer free-energy value of 4-7 kJ/mol (Figure 6A). To investigate the dynamics of this process further, we carried out a coarse-grained simulation with acylated SER181 (LCAT-mem-acyl-CG). The simulation trajectory revealed that approximately after 1.6 μs (scaled Martini time, see Table II) a cholesterol molecule spontaneously diffused to the active site (Figure 6C and Movie S5) agreeing with our free-energy calculations. This was also registered by calculating the contacts between the side chain atoms of SER181 and the hydroxyl group of CHOL molecules (See Fig 6B). In addition, a closer inspection unveiled that the exchange of cholesterol molecules between the active site of LCAT and the lipid membrane can occur if the esterification reaction does not take place (Movie S6).

**Figure 6.**
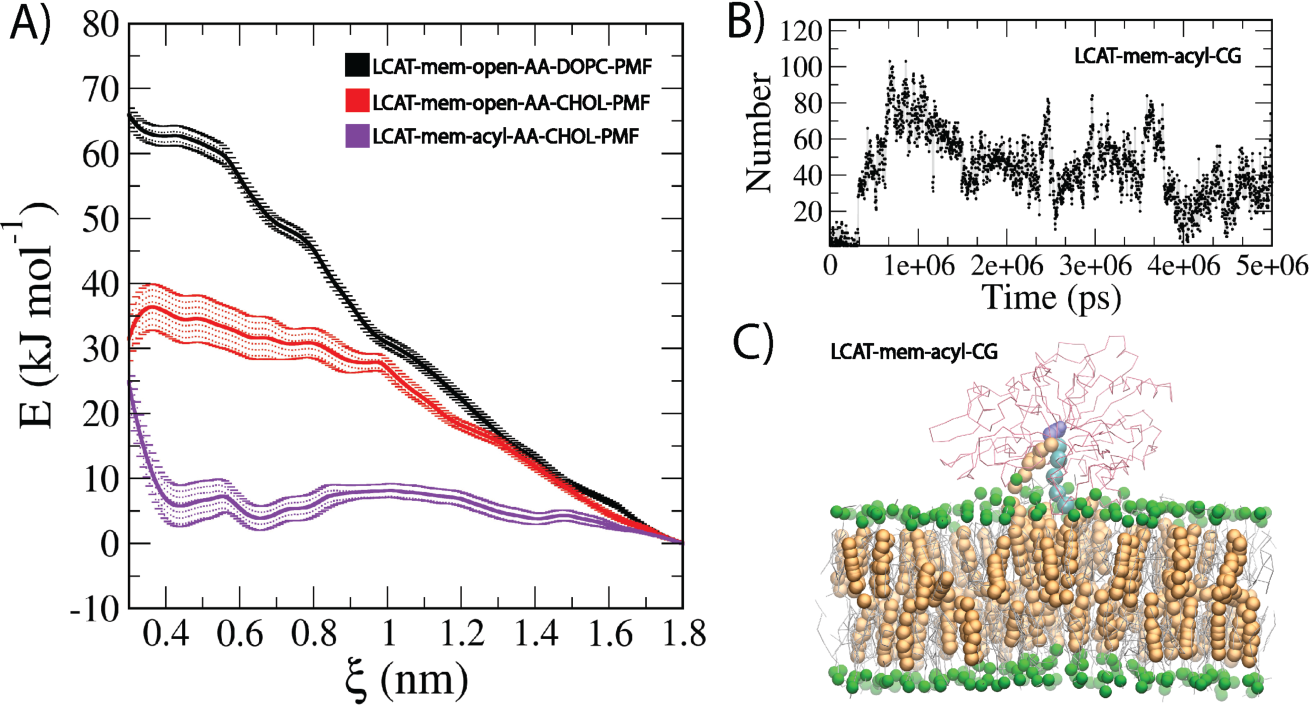
A) Potential mean force profiles (PMFs) for DOPC and CHOL calculated in LCAT-mem-open-AA system. In addition, the PMF profile for CHOL in LCAT-mem-acyl-AA systems is shown. B) The number of contacts between the oleate chain beads linked to SER181 and cholesterol beads as a function of time. C) Snapshot from LCAT-mem-acyl-CG simulation showing the location of CHOL molecule in the active site of LCAT. Cholesterol molecules are rendered as orange spheres, DOPC molecules as grey sticks, the phosphate beads of DOPC as green spheres, LCAT backbone atoms as red sticks, the oleate chain beads linked to SER181 as cyan spheres, and SER181 as violet spheres

## DISCUSSION

In the present study, we have studied the interactions of LCAT with lipid surfaces and apoA-I derived peptides to gain novel information regarding the molecular mechanism behind the LCAT catalyzed cholesterol esterification taking place at the surface of lipoprotein particles. To achieve this, we employed extensive atomistic and coarse-grained molecular dynamics simulations as well as experimental surface sensitive biophysical methods.

The simulation results showed that the lid-region of LCAT forms a coil secondary structure accompanied with relatively high residual fluctuations when compared to the other structural parts of LCAT. The fact that the lid regions are partly missing from the X-ray structures of LCAT is backing our simulation results showing a highly mobile region with no definite structural conformation (Ruwanthi N. Gunawardane et al., 2016; Manthei et al., 2017; Derek E. Piper et al., 2015). Another important finding was that the highly dynamical lid-loop shields the non-polar amino acids located at the tunnel opening from the water. Moreover, we suggest that when the lid-loop undergoes the conformational change from the closed to the open state two salt-bridges are broken and four intra-lid hydrogen bonds are formed. These findings support the view that the lid functions as a dynamical gate regulating the access of phospholipids and cholesterol molecules into the active site. In general, lipases are known to possess open and closed states (Glukhova et al., 2015; Mead et al., 2002; Yang and Lowe, 2000). For example, the X-ray structures of pancreatic lipase indicate that the opening of the active site is accompanied with considerable structural changes of the lid region governed by non-polar contacts, hydrogen bonds and salt bridges (Van Tilbeurgh et al., 1993). Although we have studied a structurally different enzyme, these findings are consistent with our results. Based on the current findings it is tempting to speculate that the closed state of LCAT is stabilized in water by the lid-mediated shielding of the non-polar amino acids at the active site tunnel opening and the presence of two salt-bridges. It is possible, therefore, that the entropic cost associated with conformational switching from the closed to open state in water, mainly arises from exposing the lid-covered non-polar amino acids to aqueous surroundings. After LCAT becomes bound to lipoprotein particles, presumably this entropic cost decreases since the non-polar residues at the tunnel opening may become buried in a hydrophobic lipid matrix as shown by our atomistic and CG-simulations. Thus, it is possible to hypothesize that this could be one of the reasons why the lid-open state could be favored when LCAT is bound to lipoprotein particles. Yet, CG-simulations revealed that LCAT can strongly interact with lipids even in the lid-closed state which was accompanied by a greater distance between the α/β hydrolase domain and the lipid bilayer surface. This conformation does not allow the burial of the tunnel opening non-polar amino acids in lipids, which we propose to be a prerequisite for the entry of lipids to the active site.

Interestingly, according to our simulations, CHOL molecules prefer to accumulate next to the membrane penetrating region of LCAT. This finding may be due to packing defects between LCAT and phospholipids. The tendency of CHOL molecules to concentrate adjacent to the tunnel opening non-polar amino acids indicates its important role in regulating the accessibility of CHOL molecules into the active site over phospholipids. This could be one of the reasons why LCAT is the sole enzyme among lipases catalyzing the esterification of CHOL molecules. This issue can be further clarified by examining the lipid surface interaction of other lipases.

Our QCM measurements revealed that the peptides derived from the LCAT-interaction region of apoA-I increased the binding of LCAT to lipid surfaces, while the control peptide 185-205 and the mutant peptide 150-170-Y166F did not facilitate the LCAT interaction with lipids. These findings agree with previous experimental reports showing that apoA-I increases the binding of LCAT to lipids compared to apolipoprotein-free small unilamellar vesicles (Jin et al., 1999; Jonas, 2000). Earlier it has been shown that the apoA-I mutation Y166F decreases the surface binding and activity of LCAT which is in agreement with our experimental results (Gu et al., 2016; Wu et al., 2007). In general, many studies have pointed out that the charged and polar amino acids located in the central parts of apoA-I are important in LCAT activity (Sorci-Thomas et al., 2009; Sorci-Thomas and Thomas, 2002). One of our unanticipated finding was that although some of the peptides did not interact with lipids alone, they still contributed strongly to the lipid attachment of LCAT. Namely, peptides 135-155 and 150-170 did not interact with lipids at all, and peptide 122-142 showed only a modest interfacial activity. Nevertheless, these peptides greatly increased the binding of LCAT to lipid surfaces when compared to the more surface-active peptides 150-170-Y166F and 185-205. Interestingly, the computational free-energy calculations revealed that all the peptides studied must adopt fully or partial amphipathic alpha-helical conformations before they can interact with lipid surfaces. It can thus be suggested that the LCAT-activating central region of apoA-I may adopt water exposed loop conformations before interacting with LCAT. The interaction of loop structures with LCAT could be followed by a conformational change from the coil to a-helix leading to stronger binding of LCAT to HDL particles. The MP-SPR results support this view since it was shown that the apoA-I derived peptides can bind to LCAT without the presence of a lipid surface. Further, based on the free-energy calculations of LCAT, it is evident that apoA-I derived peptides do not increase the binding of LCAT-peptide complexes to lipid surfaces by merely changing the lid from closed- to the open-state since the lipid binding free-energy for the open state of LCAT was lower compared to the open state. Overall, these findings strengthen the results shown in the previous experimental studies demonstrating that the central region of apoA-I possesses much lower α-helical stabilities and, thus, increased the tendency to form looped structures compared to other regions (Chetty et al., 2013; Sevugan Chetty et al., 2012; Wu et al., 2007).

The free-energy calculations produced in the present study support the view that the entry of lipids into the active site of LCAT likely occurs without the direct involvement of apoA-I. One major finding supporting this view was that the calculated transfer free-energy of DOPC from the lipid bilayer into the active site (65 kJ/mol) is congruent with the experimentally determined activation energies of LCAT in reconstituted HDL particles with varying PL species (53-76 kJ/mol) (Jonas, 1986; Parks and Gebre, 1997). This result is also consistent with the data demonstrating that the apolipoprotein content of HDL particles does not affect the activation energy of LCAT (Jonas et al., 1987). The second major finding was that the acylation of SER181 renders the transfer free-energy of cholesterol much lower when compared to the non-acylated case (4-7 kJ/mol vs. 35 kJ/mol). With respect to this finding, we observed that CHOL molecules spontaneously diffused into the active site in CG-simulations when SER181 was acylated. The CG-simulations further revealed that the exchange of CHOL molecules in the active site took place in 20 μs. Thus, the atomistic free-energy calculations and CG-simulations showed that the diffusion of lipids, especially in the case of CHOL molecules, to the active site of LCAT is not directly assisted by apoA-I. Interestingly, a recent research study revealed that molecular agents targeted for the lipid binding site of LCAT can increase the V_max_ of LCAT (Freeman et al., 2017). Surprisingly, the compound also activated LCAT deficiency causing mutants which raises hopes for treating these disorders in the future. It was identified with molecular modelling that the LCAT activating compound A formed a hydrophobic adduct with CYS31 located in the active site. Thus, one reason for the higher LCAT activity could be that the compound A renders the active site of LCAT energetically more favorable to PLs to diffuse in. In other words, the drug lowers the activation energy of LCAT, which is essentially covered by the transfer of a PL from the lipid monolayer to the active site.

To conclude, our results suggest that the initial binding of LCAT to lipoprotein surface occurs by the help of non-polar amino acids in the membrane-binding domain. The initial lipid interaction of LCAT is likely followed by a yet unknown interaction with apoA-I which enables the opening of the lid and the conformational change of LCAT to the configuration where the tunnel opening nonpolar amino acids become buried in lipids. In addition, free cholesterol molecules are attracted adjacent to the tunnel opening. Meanwhile, the amino acids 135-170 of apoA-I increases the binding of LCAT to lipoproteins. Thus, apoA-I enables the diffusion of PLs to the active site via conformational and structural changes which leads to the formation of the acyl-intermediate of LCAT. This, in turn, greatly facilitates the diffusion of CHOL molecules to the active site without the direct involvement of apoA-I. The evidence from this study suggests that the role of apoA-I and other apolipoproteins in activating LCAT is to adjust the conformation of LCAT with respect to the lipid surface, increase the binding of LCAT and drive the lid to the open state. The reasons for varying LCAT activation potencies of different apolipoproteins could be speculated to arise from their different capacity to control these features (Jonas, 1986). What is highly surprising is that the apoA-I derived peptides can increase LCAT binding to lipids, even if some of them do not interact with lipids individually. These findings encourage further investigations and modifications of these peptides aiming to increase the activity of LCAT. Eventually, this approach may i.e. lead to the development of more advanced apolipoprotein mimetic peptides that could be used in the treatment of different metabolic disorders such as dyslipidaemia and LCAT deficiencies (Navab et al., 2015, 2010). Overall, the current research study provides a novel framework for the exploration of LCAT activation by apolipoproteins, apolipoprotein mimetic peptides, and other pharmacological compounds.

## MATERIALS AND METHODS

### Computational procedures

#### Construction of simulation systems

The high-resolution X-ray structures of LCAT were acquired from the Brookhaven databank with the accession codes of 4XWG and 5BV7 (Ruwanthi N Gunawardane et al., 2016; Derek E. Piper et al.,2014). In addition, all the mutated residues (C31Y in 4XWG; L4F and N5D in 5BV7) were changed back to ones matching with the native LCAT. The 4XWG structure is considered to be the closed and 5BV7 structure the open conformation of LCAT based on the lid-loop orientation with respect to the active site tunnel opening. The Modeller program was utilized to construct the missing lid-loop regions in both structures (Webb and Sali, 2014). The default modelling parameters were utilized and the structures with the highest DOPE scores were chosen in both cases for further studies.

Two LCAT-in-water systems were constructed: one for the open and one for the closed lid-loop conformation. In the rest of the article, we refer to these systems by abbreviations LCAT-water-open and LCAT-water-closed. The dimensions of LCAT-water-open and LCAT-water-closed simulation boxes were 8.5x7x7 nm. The open and closed LCAT enzymes were solvated with 12 000 water molecules. In addition, nine sodium ions were added to neutralize the simulation systems. In the construction of LCAT-lipid bilayer systems, we used a pre-equilibrated DOPC/CHOL bilayer consisting available in the website of the Slipid developers. To construct atomistic lipid membrane systems, the LCAT was placed on the surface of DOPC/CHOL bilayer so that the non-polar residues of the tunnel-opening and the membrane binding domain were buried in the lipid matrix and the active site tunnel opening was pointing towards the lipid bilayer. This was done for both the lid-open and –closed LCAT structures. The DOPC and CHOL molecules overlapping with LCAT were removed from the systems. Following this, the systems were solvated approximately with 20 000 water molecules and nine sodium ions were added to neutralize the systems. One additional atomistic system was constructed for the open LCAT lid-loop conformation where SER181 was acylated with an oleic acid. We refer to these all-atom lipid membrane models by LCAT-mem-open-AA, LCAT-mem-closed-AA, and LCAT-mem-acyl-AA abbreviations. The starting structures for the corresponding coarse-grained LCAT-lipid systems representing these three atom-scale models were constructed similarly (LCAT-mem-open-CG, LCAT-mem-closed-CG, and LCAT-mem-acyl-CG). In Table 2, a more detailed list of molecules in each simulation system is presented.

**Table 2.**
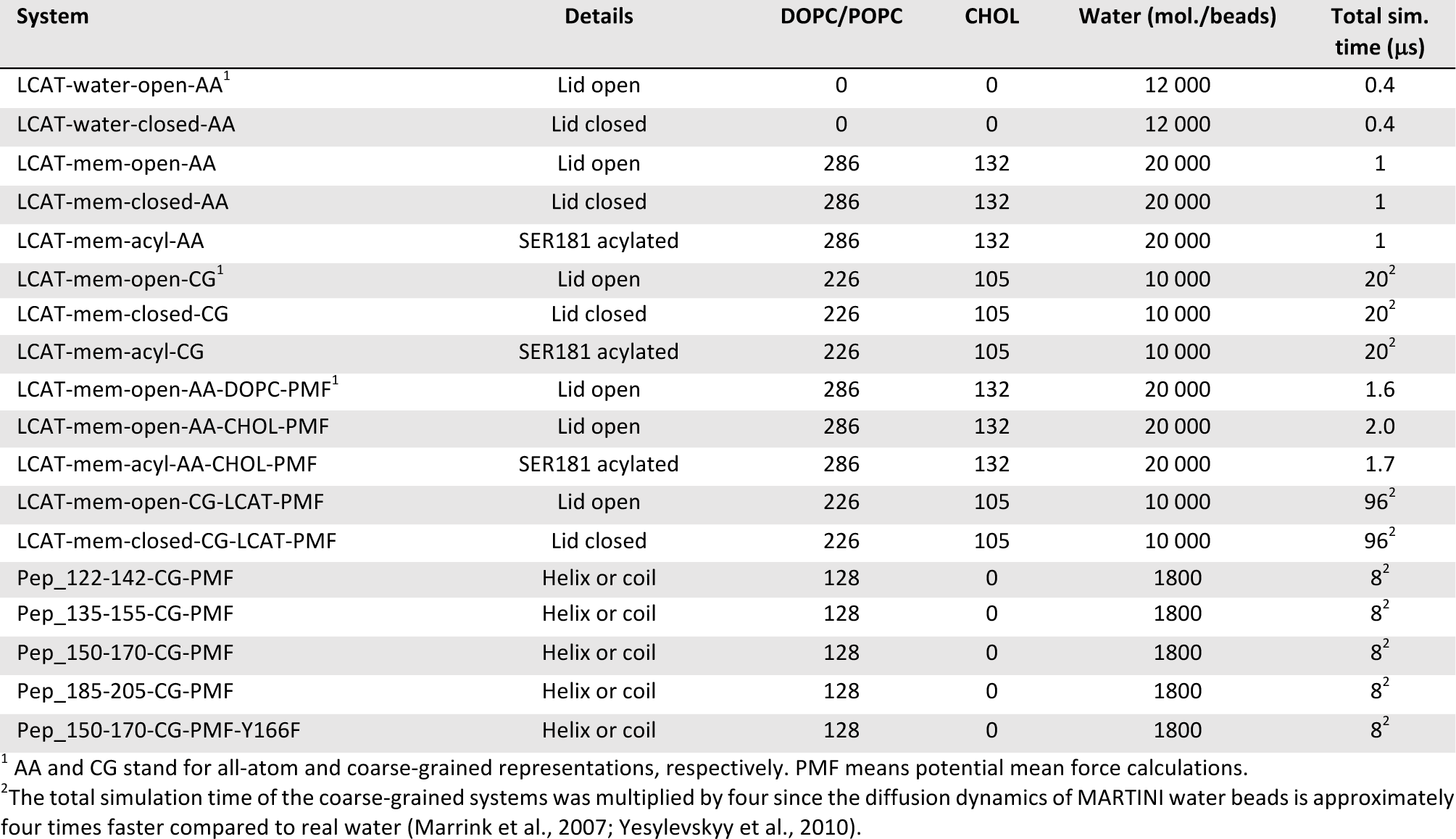
Simulated systems with their molecular compositions and simulation times.

| System | Details | DOPC/POPC | CHOL | Water (mol./beads) | Total sim. time (μs) |
| --- | --- | --- | --- | --- | --- |
| LCAT-water-open-AA<sup>1</sup> | Lid open | 0 | 0 | 12 000 | 0.4 |
| LCAT-water-closed-AA | Lid closed | 0 | 0 | 12 000 | 0.4 |
| LCAT-mem-open-AA | Lid open | 286 | 132 | 20 000 | 1 |
| LCAT-mem-closed-AA | Lid closed | 286 | 132 | 20 000 | 1 |
| LCAT-mem-acyl-AA | SER181 acylated | 286 | 132 | 20 000 | 1 |
| LCAT-mem-open-CG<sup>1</sup> | Lid open | 226 | 105 | 10 000 | 20<sup>2</sup> |
| LCAT-mem-closed-CG | Lid closed | 226 | 105 | 10 000 | 20<sup>2</sup> |
| LCAT-mem-acyl-CG | SER181 acylated | 226 | 105 | 10 000 | 20<sup>2</sup> |
| LCAT-mem-open-AA-DOPC-PMF<sup>1</sup> | Lid open | 286 | 132 | 20 000 | 1.6 |
| LCAT-mem-open-AA-CHOL-PMF | Lid open | 286 | 132 | 20 000 | 2.0 |
| LCAT-mem-acyl-AA-CHOL-PMF | SER181 acylated | 286 | 132 | 20 000 | 1.7 |
| LCAT-mem-open-CG-LCAT-PMF | Lid open | 226 | 105 | 10 000 | 96<sup>2</sup> |
| LCAT-mem-closed-CG-LCAT-PMF | Lid closed | 226 | 105 | 10 000 | 96<sup>2</sup> |
| Pep_122-142-CG-PMF | Helix or coil | 128 | 0 | 1800 | 8<sup>2</sup> |
| Pep_135-155-CG-PMF | Helix or coil | 128 | 0 | 1800 | 8<sup>2</sup> |
| Pep_150-170-CG-PMF | Helix or coil | 128 | 0 | 1800 | 8<sup>2</sup> |
| Pep_185-205-CG-PMF | Helix or coil | 128 | 0 | 1800 | 8<sup>2</sup> |
| Pep_150-170-CG-PMF-Y166F | Helix or coil | 128 | 0 | 1800 | 8<sup>2</sup> |

<sup>1</sup> AA and CG stand for all-atom and coarse-grained representations, respectively. PMF means potential mean force calculations.
<sup>2</sup> The total simulation time of the coarse-grained systems was multiplied by four since the diffusion dynamics of MARTINI water beads is approximately four times faster compared to real water (Marrink et al., 2007; Yesylevsky et al., 2010).

#### Simulation force fields and parameters

Molecular dynamics simulations were carried out with the GROMACS simulations package (version 5.1.2) (Berendsen et al., 1995; Hess et al., 2008). The Slipids and AMBER99SB-ILDN force fields were utilized for lipids and protein molecules in atomistic simulations, respectively (Jämbeck and Lyubartsev, 2012; Lindorff-Larsen et al., 2010). The MARTINI force field was used for the coarsegrained (CG) representations (Marrink et al., 2007; Monticelli et al., 2008). The polarizable water and protein models of the MARTINI force field were utilized (De Jong et al., 2013; Yesylevskyy et al., 2010).

Concerning the atomistic simulations, all systems were first energy minimized with the steepest descent algorithm. After this, short 10 ns equilibrium simulations were carried out to stabilize pressure fluctuations that were followed by 400 ns (LCAT-water systems) or one-microsecond production simulations (LCAT-mem systems). The time step was set to 2 fs, and the temperature and pressure were maintained at 310 K and 1 bar, respectively. The Berendsen coupling schemes were utilized for achieving constant temperature and pressure during the short equilibrium period of atomistic systems (Berendsen et al., 1984). After this the Nose-Hoover and the Parrinello-Rahman coupling algorithms were employed for treating temperature and pressure in the atomistic systems, respectively (Evans and Holian, 1985; Parrinello and Rahman, 1981). The isotropic or semi-isotropic pressure coupling scheme was used in LCAT-water and LCAT-mem systems, respectively. Water plus ions, lipids, and protein were separately coupled to heat paths. For the Lennard-Jones interactions a cut-off of 1 nm was used, and the electrostatic interactions were handled with the particle-mesh Ewald method with a real space cut-off of 1.0 (Essmann et al.,1995). The LINCS algorithm was applied to constrain covalent hydrogen bond lengths (Hess et al., 1997).

Regarding the coarse-grained simulations, the temperature and pressure were handled with the v-rescale and the Parrinello-Rahman schemes, respectively (Bussi et al., 2007; Parrinello and Rahman, 1981). Coupling constants of 1 ps^-1^ and 12 ps^-1^ were used for the temperature and pressure schemes, respectively. Lipids, protein and water molecules were separately coupled to a heat path. The reaction-field field electrostatics and Lennard-Jones interactions were employed with cut-offs of 1.1 nm. The relative electrostatic screening was set to 2.5 since the polarizable water and protein models of MARTINI force field were used. Time step of 20 fs was used and all coarse-grained simulations were simulated up to 20 us (scaled Martini time). The ElNeDyn scheme was employed for LCAT (Periole et al., 2009).

#### Free-energy calculations

For calculating the free-energy profiles for PC or CHOL molecules entering the active site of LCAT, either a CHOL or a PC molecule was pulled from the lipid bilayer to the active site of LCAT in the lid-open state to generate umbrella windows (these systems are coined as LCAT-mem-open-DOPC-PMF-AA and LCAT-mem-open-CHOL-PMF-AA). The free-energy profiles for PC and CHOL molecules were calculated utilizing atomistic simulations. In addition, the free-energy profile for a CHOL molecule calculated in the LCAT-mem systems where SER181 was acylated (LCAT-mem-acyl-CHOL-PMF-AA). The oxygen atom of CHOL or phosphorous atom of DOPC was pulled towards the center of mass of SER181 residue with a pull rate of 0.001 nm/ps and a force constant of 10 000 kJ/mol·nm^2^. The resulting reaction coordinate was divided into 39 umbrella windows. The force-constant of the spring was set to 2000 kJ/mol·nm^2^. Each umbrella window was sampled up to 40-50 ns, thus the total simulation time ranged from 1.6 to 2.0 μs per system. The last 34-28 ns of the simulation trajectories were used to construct the PMF profiles.

For determining the lipid-binding free-energies for different apoA-I derived peptides at the coarse-grained level, each peptide was pulled from the water phase to the center of POPC bilayer with a pull rate of 0.001 nm/ps and a force-constant of 20 000 kJ/mol·nm^2^. This was followed by a generation of 20 umbrella windows along the bilayer normal. Each window was sampled up to 100 ns (400 ns in scaled MARTINI time) and the centers of mass of peptides were constrained to the center of each umbrella window by using a force constant of 500 kJ/mol·nm^2^. The last 320 ns of the simulation trajectories were used to construct the PMF profiles. To estimate the effect of secondary structure to the binding of free-energies, potential mean force profiles were calculated for peptides fully adopting alpha-helical or coil secondary structures.

In addition to the above systems, also the free-energy profiles for LCAT along the lipid membrane normal was estimated. First, LCAT was pulled from the surface of DOPC-CHOL membrane to the water phase by using a constant pulling speed of 0.0002 nm/ps and a force constant of 20 000 kJ/mol·nm^2^. The pulling was done for end structures of LCAT-mem-open-CG and LCAT-mem-closed-CG simulations. Next, the reaction coordinate was divided into 30 different umbrella windows. Each window was sampled up to 0.8 μs (3.2 μs in scaled MARTINI time) in with a force constant of 1000 kJ/ mol·nm^2^. Thus, the total simulation time for each LCAT system was 96 μs (scaled Martini time). The last 60 μs of the simulation trajectories were used in the construction of the PMF profiles.

#### Analysis methods

All the analysis programs reported here are part of the GROMACS simulations package unless mentioned otherwise. The gmx rmsf and gmx rmsd programs were used to produce root mean square fluctuation (RMSF) and deviation (RMSD) graphs. Only the backbone atoms of LCAT were included in the RMSD and RMSF analysis. The RMSF profiles were calculated as a function of amino acid residues of LCAT after the RMSD profiles were stabilized (after 100 ns). The secondary structure of LCAT was monitored as a function of time with the secondary structure plugin of the VMD (Humphrey et al., 1996). The average number of hydrogen bonds was calculated utilizing the gmx hbond analysis tool. The maximum distance between a donor and an acceptor was set to 0.35nm, and the maximum angle formed by hydrogen bonding atoms hydrogen-donor-acceptor was set to 30 degrees. The number of salt-bridges in the structure of LCAT concerning the lid-region was calculated by the gmx mindist program with a distance criterion of 0.6 nm between the charged side chain central carbon atoms of negatively and positively charged amino acids (GLU-ARG, GLU-LYS, ASP-ARG, and ASP-LYS). The solvent accessible surface areas (SASA) for the selected amino acids were calculated using the gmx sasa program. The distance and tilt analysis were carried out with the gmx mindist and gmx gangle programs. These analyses were conducted after the number of contacts between protein and head group atoms of DOPC were in equilibrium. For producing 2D-spatial density maps the gmx densmap program was used. The XY-planes of simulation systems were divided into 100x100 or 36x36 square bins, in which the number of CHOL atoms was calculated in each time frame and summed over the whole simulation time to produce number density profiles. Before the calculation, the membrane puncturing region of LCAT (the center of mass of residue PHE67) was placed to the center of the simulation box. The gmx_wham program was utilized to derive the potential mean force profiles from the umbrella sampling simulations. The bootstrapping method with 50 bins was used to estimate the errors for the PMF profiles. All visualizations in the article were made either with the Python codes or the VMD visualization package (Humphrey et al., 1996).

### Experimental procedures

#### Materials

POPC (1-palmitoyl-2-oleoyl-*sn*-glycero-3-phosphocholine), dissolved in chloroform (25 mg/mL) was obtained from Avanti Polar Lipids (Alabaster, USA). Calcium chloride, 4-(2-hydroxyethyl)-1-piperazineethanesulfonic acid (HEPES), 3-[(3-)cholamidopropyl)dimethylammonio]-1-propanesulfonate (CHAPS), sodium chloride, ethylene diaminetetraacetic acid (EDTA), sodium azide, and potassium dihydrogen phosphate were obtained from Merck Sigma-Aldrich (Darmstadt, Germany). Potassium chloride was obtained from Honeywell Riedel de Haën (Seelz, Germany), disodium hydrogen phosphate was obtained from Fisher Scientific (Hampton, USA). Peptides and N-terminal biotin-labeled peptides were synthesized by Peptide Protein Research Ltd, Hampshire, U.K. and had the following amino acid sequences: (i) 122 - 142: LRAELQEGARQKLHELQEKLS (ii) 135 - 155: HELQEKLSPLGEEMRDRARAH (iii) 150 - 170: DRARAHVDALRTHLAPYSDEL (iv) 150 - 170-Y166F: DRARAHVDALRTHLAPFSDEL (v) 185 - 205: GGARLAEYHAKATEHLSTLSE. Lyophilized, glycosylated human LCAT protein (Sino Biological Inc., Beijing, China) was reconstituted with 400 μl of ultrapure, sterile water to yield LCAT at 5 μM in PBS (manufacturer specifications). Ultra-pure water used for preparation of the buffer and in all measurements, were prepared with a Milli-Q purification system, having a resistivity of 18 MΩ·cm and TOC level of < 5 ppm. The following buffers were used: 20 mM phosphate buffer saline (PBS), 1 mM EDTA, 1 mM Na Azide (pH 7.4; buffer A); 20 mM HEPES 150 mM NaCl (pH 7.4; buffer B).

#### The multi-parametric surface plasmon resonance measurements

Measurements were performed with a multi-parameter SPR Navi™ 200 (BioNavis Ltd., Tampere, Finland) instrument. The setup was equipped with two incident laser wavelengths, 670 nm and 785 nm, two independent flow channels, inlet tubing and outlet (waste) tubing, and an autosampler. Both of the flow channels were measured simultaneously with 670 nm and 785 nm incident light. The measurement temperature was kept constant at 20 °C. Biotin coated SPR sensors (Bionavis Ltd., Tampere, Finland) were placed in the flow-cell. The flow rates used for the streptavidin interaction with biotin, biotin-peptides with streptavidin, and for LCAT interaction with immobilized peptides were 10 μL/min. SPR spectra were recorded after introduction of buffer A into the flow-cell for 10 – 30 min until a stable baseline was achieved after which the surface was fully saturated with streptavidin during a 10 minutes serial injection at 1.9 μM to saturate the SPR sensor. It was observed that after 5 minutes the SPR sensor was fully coated with streptavidin, therefore the method was adapted to 5 minutes. In the second phase of the measurement, in DMSO solubilized biotin-peptides with a final concentration of 100 μM were injected in parallel into both flow channels for 7 minutes at 10 μM to cover the SPR sensor fully, i.e. until all streptavidin bindings side were fully occupied. Parallel injections ensured that 2 peptides were analyzed simultaneously. Increasing concentrations of LCAT were then injected in both channels via serial injection, to ensure the lowest variation of LCAT injection for 15 minutes at increasing concentrations (156, 313, 625, 1250, 2500, and 5000 nM) to determine the binding constant of the LCAT-peptide complexes. The data was fitted in OriginPro (v. 8.6, OriginLab Corp., Northampton, MA, USA) according to the Langmuir model to obtain the dissociation constants (K_D_) and the responses of saturated binding (R_max_):

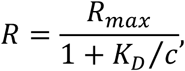

where R is the final response after each injection and c is the concentration of LCAT in each injection.

#### The quartz crystal microbalance measurements

Prior to the QCM measurements, the PBS buffer formed after reconstitution LCAT was exchanged for buffer A due to the extreme high sensitivity of the QCM method to slight differences in ionic strength of the background buffer. The LCAT solution was placed in an Amicon Ultra-0.5 10K Centrifugal Filter Device (Merck Millipore Ltd, Tullagreen Carrigtohill, Ireland) and equal volumes of buffer A were filtered through ten times at 14 000 x g, after which 400 μl of buffer A was added to fully recover LCAT at 5 μM. The peptides were dissolved into buffer A at a final concentration of 100 μM.

An impedance-based quartz crystal microbalance instrument (KSV Instruments Ltd, Helsinki, Finland) was used for the measurements. The measurement temperature was kept constant at 20 °C. Before measurements, silica-coated QCM sensors (Q-Sense Inc./BiolinScientific, Västra Frölunda, Sweden) were first flushed with 70 % ethanol and ultrapure water, dried under nitrogen flow, and finally oxygen plasma-treated (PDC-002, Harrick Plasma, Ithaca, USA) for 5 minutes at 29.6 W and 133-173 Pa. Samples were injected into the measurement chamber with a peristaltic pump system (Ismatec/Cole-Parmer GmbH, Wertheim, Germany). Small unilamellar POPC liposome vesicles (SUVs) which were used to form supported lipid bilayers inside the QCM measurement chamber were made using thin film hydration method followed by extrusion (11 times) through a 50-nm polycarbonate filter membrane at 60 °C. The resulting SUVs (10 mg/mL in buffer B) had a mean number average particle size of 65 ± 7 nm and polydispersity index of 0.167 ± 0.008 (determined from three individual measurements by Zetasizer APS instrument, Malvern Instruments Ltd, Worcestershire, UK).

After the baseline of the signal at different frequency overtones (3, 5 and7) was stabilized with buffer B, supported lipid bilayers (SLBs) were formed by flowing liposome solution (0.15 mg/mL in buffer B + 5 mM CaCl_2_ at 250 μL/min) over the crystal surface for 5 minutes. Afterwards, buffer A was injected, and the signal was left to stabilize. An additional injection of ultrapure water was added to ensure the complete formation of an SLB and absence of intact vesicles on the measurement surface. The overlap of the changes in normalized overtone frequencies (-26 Hz) was considered as a confirmation of the presence of a good-quality SLB. The flow speed was reduced to 50 μL/min and the measurements were performed by injecting peptides at 100 μM in buffer A, LCAT at 37.5 nM in buffer A, or combination of LCAT and peptide through the measurement chamber for a duration of 10 minutes. Between each measurement, sensors were cleaned *in situ* by sequential 2 minute injections of 20 mM CHAPS, 2 % Hellmanex, 70 % ethanol, and ultrapure H_2_O.

Data analysis was performed using Origin Pro (v. 8.6, OriginLab Corp., Northampton, MA, USA). For each measurement, frequency overtone signals (3, 5 and 7) were normalized, averaged and baseline corrected. Neither LCAT nor peptides induced changes in viscoelastic properties of the bilayer, which was seen as negligible changes in the recorded energy dissipation.

## ACKNOWLEDGEMENTS

Academy of Finland is thanked for providing financial support for MC (key project funding, project no. 303884), AK (postdoctoral grant, project no. 298863) and TV (project no. 137053 and 263861). PP acknowledges a grant from the Finnish Pharmaceutical Society. CSC – IT Centre for Science Ltd. (Finland) is acknowledged for computational resources. This article is dedicated to the memory of Dr. Marja T. Hyvönen.

